# Integrity of corpus callosum is essential for the cross-hemispheric propagation of sleep slow waves: a high-density EEG study in split-brain patients

**DOI:** 10.1101/756676

**Authors:** Giulia Avvenuti, Giacomo Handjaras, Monica Betta, Jacinthe Cataldi, Laura Sophie Imperatori, Simona Lattanzi, Brady A. Riedner, Pietro Pietrini, Emiliano Ricciardi, Giulio Tononi, Francesca Siclari, Gabriele Polonara, Mara Fabri, Mauro Silvestrini, Michele Bellesi, Giulio Bernardi

**Author notes:** Equal contribution (Michele Bellesi, Giulio Bernardi). **Correspondence**: Giulio Bernardi, MoMiLab Research Unit - IMT School for Advanced Studies, Piazza San Francesco 19, Lucca, 55100, Italy, Michele Bellesi, School of Physiology, Pharmacology & Neuroscience - University of Bristol, Biomedical Sciences Building, University Walk, Bristol BS8 1TD, UK.

## Abstract

The slow waves of NREM-sleep (0.5-4Hz) reflect experience-dependent plasticity and play a direct role in the restorative functions of sleep. Importantly, slow waves behave as traveling waves and their propagation is assumed to reflect the structural properties of white matter connections. Based on this assumption, the corpus callosum (CC) may represent the main responsible for cross-hemispheric slow wave propagation. To verify this hypothesis, here we studied a group of patients who underwent total callosotomy due to drug-resistant epilepsy. Overnight high-density (hd)-EEG recordings (256 electrodes) were performed in five totally callosotomized in-patients (CP; 40-53y, 2F), in three control non-callosotomized neurological in-patients (NP; 44-66y, 2F, 1M epileptic), and in an additional sample of 24 healthy adult subjects (HS; 20-47y, 13F). Data were inspected to select NREM-sleep epochs and artefactual or non-physiological activity was rejected. Slow waves were detected using an automated algorithm and their properties and propagation patterns were computed. For each slow wave parameter and for each patient, the relative z-score and the corresponding p-value were calculated with respect to the distribution represented by the HS-group. Group differences were considered significant only when a Bonferroni corrected *P* < 0.05 was observed in all the CP and in none of the NP. A regression-based adjustment was used to exclude potential confounding effects of age. Slow wave density, amplitude, slope and propagation speed did not differ across CP and HS. In all CP slow waves displayed a significantly reduced probability of cross-hemispheric propagation and a stronger inter-hemispheric asymmetry. Moreover, we found that the incidence of large slow waves tended to differ across hemispheres within individual NREM epochs, with a relative predominance of the right over the left hemisphere in both CP and HS. The absolute magnitude of this inter-hemispheric difference was significantly greater in CP relative to HS. This effect did not depend on differences in slow wave origin within each hemisphere across groups. Present results indicate that the integrity of the CC is essential for the cross-hemispheric traveling of sleep slow waves, supporting the assumption of a direct relationship between white matter structural integrity and cross-hemispheric slow wave propagation. Our findings also imply a prominent role of cortico-cortical connections, rather than cortico-subcortico-cortical loops, in slow wave cross-hemispheric synchronization. Finally, this data indicate that the lack of the CC does not lead to differences in sleep depth, in terms of slow wave generation/origin, across brain hemispheres.

## Introduction

The transition from wakefulness to sleep is marked by profound changes in brain EEG activity, with a shift from the low-amplitude, high-frequency signals recorded in wakefulness to the high-amplitude, low-frequency slow waves (0.5-4 Hz) of NREM-sleep. In particular, sleep slow wave represents the EEG signature of a slow oscillation in membrane potential at neuronal level, characterized by an alternation between a hyperpolarized “silent” phase (*down-state*) and a depolarized phase of intense firing activity (*up-state*) (Steriade *et al.*, 2001). Crucially, the amount of slow wave activity (SWA, expressed as the 0.5-4 Hz EEG-signal power in NREM-sleep) represents a reliable marker of homeostatically regulated sleep need (Achermann and Borbély, 2003) and has been shown to be locally modulated in a use-dependent manner, thus implying a possible relationship with plasticity-related processes (Tononi and Cirelli, 2014). Indeed, experimental studies and computer simulations have demonstrated that not only SWA reflects experience-dependent changes in regional synaptic density/strength, but also have indicated that slow waves may play a direct role in cellular and systems restoration and in the consolidation of newly acquired memories (Tononi and Cirelli, 2014). Recent evidence also suggested a possible implication of sleep slow waves in the clearance of neurotoxic metabolic products that accumulate during wakefulness (Xie *et al.*, 2013; Hablitz *et al.*, 2019).

The sleep slow waves are not stationary events. Instead, they typically behave as traveling waves at the macro-scale level of the scalp EEG, with variable cortical origin and propagation pattern (Massimini *et al.*, 2004; Murphy *et al.*, 2009). Such a propagation is commonly assumed to reflect the structural properties of cortico-cortical white matter connections. In line with this, structural white matter properties have been found to correlate with parameters reflecting slow waves synchronization (Murphy *et al.*, 2009; Buchmann *et al.*, 2011; Piantoni *et al.*, 2013). In this perspective, the corpus callosum (CC) would be expected to represent the main route responsible for cross-hemispheric slow wave propagation. However, correlational studies and research in human models of inter-hemispheric disconnection produced contradictory findings. For instance, two studies found a positive significant correlation between macro (volume) and micro (axial diffusivity) structural properties of the CC and parameters reflecting slow wave synchronization (i.e., amplitude and slope) in healthy adult individuals (Buchmann *et al.*, 2011; Piantoni *et al.*, 2013). In contrast, a more recent work failed to replicate the correlation between slow-wave slope and axial diffusivity in healthy adult subjects, and rather described a positive correlation between indices reflecting white matter damage and slow wave synchronization in patients with traumatic brain injury (TBI; Sanchez *et al.*, 2019). In addition, while studies performed in patients with agenesis of the CC (Kuks *et al.*, 1985; Nielsen *et al.*, 1993) or in epileptic patients who underwent partial or total callosotomy (Montplaisir *et al.*, 1990) showed a decreased inter-hemispheric coherence within the delta range (<4 Hz) during NREM-sleep, callosotomized patients continue to present a clear increase in inter-hemispheric coherence from wakefulness to sleep (Corsi-Cabrera *et al.*, 2006; also see Supplementary Table S1).

Given the above considerations, it is still unclear how the agenesis or the complete resection of the corpus callosum may affect sleep slow wave propagation in humans. Crucially, this matter has more general implications for the hypothesized relationship between brain structural connectivity and slow wave propagation (Murphy *et al.*, 2009), as well as for the understanding of the mechanisms that regulate slow wave synchronization in relation to plastic and developmental processes (e.g., Mascetti *et al.*, 2013; Kurth *et al.*, 2017). Moreover, the contradictory findings reported in the literature likely result mostly from methodological limitations and discrepancies. Therefore, to determine the role of inter-hemispheric white matter connections in slow wave generation and propagation, here we analyzed for the first time overnight high-density (hd-)EEG recordings collected in a sample of epileptic patients who underwent total callosotomy (complete split-brain) and in control subjects with an intact CC. In order to overcome limitations of previous studies related to the use of indirect indices of slow wave synchronization and propagation, we used validated algorithms to detect individual slow waves and to determine their specific origin and traveling pattern.

## Materials and methods

### Participants

Overnight hd-EEG recordings (256 electrodes; EGI-Philips) were performed at the Neurological Unit of the Marche Polytechnic University (Ancona, Italy) in five epileptic in-patients who underwent a total resection of the CC (CP, callosotomized patients; age range 40-53, two females; Supplementary Figure S1). Three non-callosotomized neurological in-patients (NP, non-callosotomized patients; age range 44-66, two females) were also studied under the same experimental conditions. One of these patients was diagnosed with symptomatic generalized epilepsy due to viral meningoencephalitis occurred in infancy (this subject, indicated as NP03, was marked using a distinctive color in figures). All the non-callosotomized patients have no diagnoses of any other comorbidities affecting brain function at the time of the study. Table 1 and Supplementary Table S2 report demographic and clinical characteristics for all patients. An additional control group of 24 healthy adult volunteers (HS, healthy subjects; age range 20-47, 13 females) was studied with the same hd-EEG recording system at the Lausanne University Hospital, Switzerland (analyses of NREM-sleep data from these subjects, not involving the study of inter-hemispheric slow wave propagation, have been reported in previous work; Siclari *et al.*, 2018, Bernardi *et al.*, 2019*a,b*). Prior to their inclusion into the study, HS group individuals underwent a clinical interview to exclude a history of sleep, medical and psychiatric disorders. None of the HS subjects was taking any medication at the time of the study. The study procedures were conducted under clinical research protocols approved by the local ethical committees and in accordance with the guidelines of the Declaration of Helsinki. Written informed consent was obtained from all participants.

**Table 1.** Demographic and clinical characteristics. For patients in CP and NP group demographic characteristics, questionnaires and sleep macro-architecture are presented separately for each subject. For CP patients the resection of the corpus callosum occurred in two distinct surgical interventions, for each of which the age of the patients is reported. For the HS group the values are reported as group mean ± standard deviation. Sleep stage percentages are expressed with respect to total sleep time (TST). PSQI = Pittsburg Sleep Quality Index; ESS = Epworth Sleepiness Scale; HOQ = Horne-Őstberg Questionnaire; EHI = Edinburgh Handedness Inventory; PSG = polysomnography; WASO = wake after sleep onset; CP = callosotomized patients; NP = non-callosotomized patients, HS = healthy subjects.

|  | CP01 | CP02 | CP03 | CP04 | CP05 | NP01 | NP02 | NP03 | HS (n=24) |
| --- | --- | --- | --- | --- | --- | --- | --- | --- | --- |
| Age, years | 53 | 40 | 47 | 45 | 42 | 45 | 66 | 44 | 27 ± 6 |
| Gender | M | F | F | M | M | F | F | M | 13F |
| Age at Surgery 1, years | 30 | 16 | 25 | 14 | 18 | - | - | - | - |
| Age at Surgery 2, years | 45 | 17 | 26 | 22 | 19 | - | - | - | - |
| **Questionnaires** |  |  |  |  |  |  |  |  |  |
| PSQI | 12 | 2 | 4 | 5 | 9 | 11 | 4 | 12 | 3.1 ± 1.5 |
| ESS | 23 | 4 | 2 | 0 | 13 | 7 | 18 | 1 | 5.9 ± 2.1 |
| HOQ | 59 | 44 | 43 | 61 | 54 | 52 | 34 | 45 | 51.8 ± 6.7 |
| EHI | RH | RH | RH | RH | RH | RH | RH | RH | 20RH |
| **Sleep structure** |  |  |  |  |  |  |  |  |  |
| Sleep latency, min | 20.5 | 29.5 | 23.5 | 38.5 | 34.0 | 6.0 | 6.5 | 4.0 | 15.4 ± 16.9 |
| Total sleep time, min | 244.5 | 243.5 | 185.5 | 254.0 | 120.5 | 278.0 | 251.5 | 118.5 | 256.6 ± 30.6 |
| Sleep efficiency, % | 81.4 | 81.0 | 61.7 | 84.5 | 40.1 | 92.5 | 83.7 | 39.4 | 85.5 ± 10.2 |
| WASO, min | 36 | 15.5 | 66.0 | 4.5 | 129.5 | 11.0 | 24.5 | 157.5 | 20.0 ± 15.3 |
| N1 sleep, % | 0.8 | 7.0 | 14.8 | 9.6 | 24.5 | 3.1 | 8.5 | 19.4 | 5.1 ± 5.0 |
| NREM sleep, % | 100.0 | 88.7 | 100.0 | 100.0 | 100.0 | 72.7 | 81.7 | 78.5 | 84.8 ± 6.0 |
| REM sleep, % | 0.0 | 11.3 | 0.0 | 0.0 | 0.0 | 27.3 | 18.3 | 21.5 | 15.2 ± 6.0 |

### Data Acquisition

One overnight hd-EEG recording (500 Hz sampling rate) was obtained for each subject. All recordings were initiated at the usual bedtime of each participant and interrupted at ~7AM. Given that callosotomized patients presented relatively low sleep quality with frequent awakenings, especially in the second part of the night, we extracted and analyzed only the first 5h of each recording, starting from the time of ‘lights-off’. In order to ensure across-group comparability, analyses similarly focused only on the first 5h of data also in the healthy control subjects. Three 2-min ‘resting-state’ hd-EEG recordings (6-min in total) were also collected during relaxed wakefulness with the eyes closed before sleep and in the morning, about 40 min after awakening.

### Data Preprocessing

For all patients, recordings were band-pass filtered between 0.1 and 45 Hz. Then, overnight recordings were divided into 30-sec epochs, while wake resting-state recordings were divided into 4-sec epochs. Bad channels and epochs were identified and rejected through visual inspection in NetStation 5.3 (EGI-Philips). An Independent Component Analysis (ICA) procedure was used to reduce residual ocular, muscular and electrocardiograph artifacts (EEGLAB toolbox; Delorme and Makeig, 2004). Finally, rejected bad channels were interpolated using spherical splines. A similar procedure was used to preprocess the sleep data of healthy control subjects, as described in previous work (Bernardi *et al.*, 2019*b*). Data were filtered between 0.5 and 40 Hz prior to the analyses of signal power and slow wave parameters.

### Sleep Scoring

For scoring purposes, four electrodes were used to monitor horizontal and vertical eye movements (electrooculography, EOG), while electrodes located in the chin-cheek region were used to evaluate muscular activity (electromyography, EMG). Sleep scoring was performed over 30-sec epochs according to the criteria from the American Academy of Sleep Medicine scoring manual (Iber *et al.*, 2007). Of note, all epileptic patients (CP01-CP05 and NP03) presented altered patterns of brain activity, with bursts of spike-wave discharges, during both wakefulness and sleep. Such non-physiological activity was particularly evident in four patients (CP01-CP04) and limited the possibility to accurately recognize changes in sleep depth based on standard criteria (the remaining patient, CP05, is marked using a distinctive color in figures). For this reason, a distinction between N2- and N3-sleep was not made. Two operators took care in marking periods containing large artifacts, arousals and non-physiological activity (examples of included and excluded data for epileptic patients are shown in Supplementary Figure S2). Only slow waves detected within artifact-free NREM-sleep (N2/N3) data-segments were analyzed.

### Power Computation

A current-source-density (CSD) transform was applied to all EEG recordings using the CSD-toolbox (Kayser and Tenke, 2006). This method provides a reference-independent signal and improves spatial resolution by acting as a spatial filter. For each EEG derivation, power spectral density (PSD) estimates were computed using the Welch’s method in 4-s data segments (Hamming windows, 8 sections, 50% overlap) and integrated within the delta/SWA (0.5-4 Hz) and the beta (18-35 Hz) frequency bands. In all sleep epochs, the power computation was performed over seven 4-s segments (28-s) after excluding the first and the last second of data.

### Slow wave detection

Slow waves were detected automatically in a composite EEG-signal generated from linked-mastoid referenced channels, as previously described (Siclari *et al.*, 2014; Mensen *et al.*, 2016; Bernardi *et al.*, 2018). This method provides a unique time reference (across electrodes) for each slow wave and facilitates the detection of both local and widespread events (Mensen *et al.*, 2016). Specifically, a negative-going signal envelope was calculated by selecting the fifth most negative sample across a subset of 191 electrodes obtained by excluding channels located on the neck and face regions. This approach minimizes the risk of including in the envelope potential residual high amplitude oscillations of artifactual origin. Finally, the obtained signal-envelope was broadband filtered (0.5-40 Hz) prior to the application of a slow wave detection procedure based on half-wave zero-crossings (Riedner *et al.*, 2007; Siclari *et al.*, 2014). Only half-waves with a duration comprised between 0.25 and 1.0 s were retained for further analyses. Of note, no amplitude thresholds were applied based on previous evidence indicating that: i) slow waves with peak-to-peak amplitude < 75μV show the clear homeostatic changes commonly attributed to the slow waves of NREM-sleep (Riedner *et al.*, 2007; Bernardi *et al.*, 2018); ii) the application of an amplitude threshold may actually preferentially select a minority of very large slow waves that have been shown to display different regulation and synchronization mechanisms as compared to the majority of slow waves (Siclari *et al.*, 2014, 2018; Mensen *et al.*, 2016; Bernardi *et al.*, 2018, 2019*a*; Spiess *et al.*, 2018). For all the detected slow waves, various parameters of interest were calculated and stored for a subsequent evaluation, including negative amplitude (μV), descending slope (between the first zero-crossing and the maximum negative peak; μV/ms) and involvement (mean EEG-signal calculated for all electrodes in an 80-ms window centered on the wave peak; μV). Moreover, the slow wave density (number of waves per minute) was computed in each sleep epoch (epochs in which artifactual or non-physiological activity occupied more than the 75% of time were excluded).

### Scalp involvement distribution

For each subject, the involvement distribution (across channels) of all slow waves was analyzed using principal component analysis (PCA) as described in previous work (Bernardi *et al.*, 2018). We recently showed that in healthy adult subjects the 95% of the variance related to slow wave involvement is explained by three principal components (PCs), with maxima located in the centro-frontal area (~70% of total variance), anterior or posterior areas (~20%) and left or right hemispheres (~5%). Here we hypothesized that callosotomized patients would present an increased variance explained by the last, uni-hemispheric PC at the expenses of the other two, symmetrical components. To test this hypothesis, we first verified through visual inspection that similar PCs explaining a similar amount of total variance were present in HS (95.0 ± 1.5%, range 92.3-97.0), NP (96.6 ± 0.6%, range 96.2-97.3; relative to HS all P*unc.* > 0.099, |z| < 1.652) and CP (94.6 ± 1.3%, range 93.1-95.9; all P*unc.* > 0.193, |z| < 1.302) subjects. Then, the PCs-space of each subject was rotated into a common, reference-PCs-space using the Procrustes transformation (Schönemann, 1966; Haxby *et al.*, 2011). The Procrustes transformation is an orthogonal transformation that minimizes the Euclidean distance between two sets of paired vectors. The reference-space was selected by iteratively applying the transformation over pairs of subjects of the HS group and then identifying the coordinate system (i.e., the subject) presenting the smallest distance with respect to the coordinate systems of all tested subjects (Haxby *et al.*, 2011). Finally, the Procrustes transformation was applied to remap the original PCs-space of each subject (including the patients), into the new reference-PCs-space. This procedure allowed us to compare directly the first three PCs (and their explained variances) across individuals.

### Slow wave propagation

For each detected slow wave, the pattern of propagation was calculated as described in previous work (Mensen *et al.*, 2016; Bernardi *et al.*, 2018). In brief, we first determined for each electrode the timing of any local negative maxima that occurred within a time-window of 300 ms centered on the maximum negative peak (reference peak) of the examined slow wave. After exclusion of small local maxima using a threshold corresponding to the 10% of the largest maximum peak across all electrodes, the timing of the negative maximum closer in time to the reference peak was identified. A cross-correlation procedure (Mensen *et al.*, 2016) was applied to exclude channels in which the negative wave was excessively dissimilar from the ‘prototype’ slow wave, defined as the wave with the largest negative peak at the reference peak timing across all channels. Specifically, channels showing a maximal cross-correlation value smaller than a threshold corresponding to the 25^th^ percentile of the distribution of all cross-correlation values were discarded. The latency of all remaining local peaks was subsequently used to create a preliminary scalp ‘delay-map’. A spatio-temporal clusterization procedure was applied to exclude potential propagation gaps: local peaks of two spatial neighbor electrodes had to be separated by less than 10 ms in order to be considered as part of the same propagation cluster. Then, the propagation cluster including the prototype wave was identified and the final delay-map was extracted. The minimum delay, corresponding to the slow wave origin, was set to zero. Finally, a three-dimensional gradient (two for direction, one for timing) was computed from the delay-map to identify the main streamlines of propagation for each slow wave. Up to three streamlines were extracted: the longest displacement (the distance between the start and end points of the wave); the longest distance traveled (the cumulative sum of all coordinates of the line); and the stream of the most angular deviation from the longest displacement (minimum trajectory difference was set to 45°). The streamline corresponding to the longest distance traveled was used to compute the slow wave speed (distance divided by maximum delay; m/s).

### Cross-hemispheric propagation

Information obtained from the propagation analysis was used to compute parameters reflecting the degree of cross-hemispheric propagation of sleep slow waves. First, for each slow wave we determined whether at least one of the propagation streamlines passed the midline (nasion-inion axis). This information was used to compute the proportion of cross-hemispheric slow waves in each subject (% of all the detected slow waves). Second, we determined the relative distribution of electrodes involved in the same (propagating) slow wave across the two hemispheres. This information was used to compute an index of channel recruitment asymmetry, defined as the number of channels in the hemisphere with less involved electrodes, divided by the total number of involved channels (%). In this index, a value of 50% indicates a symmetric distribution, while a value of 0% indicates a unilateral wave. This second parameter was also computed for slow waves subdivided into five amplitude percentile classes: 0-20, 20-40, 40-60, 60-80, 80-100.

### Inter-hemispheric differences in slow wave density

The CC could be involved not only in the propagation of individual slow waves but also in homogenizing sleep depth across the two hemispheres. In other words, the lack of inter-hemispheric connections could allow for the emergence of hemispheric asymmetries in the relative density of large slow waves, potentially up to a marked uni-hemispheric sleep. To test this hypothesis, slow waves were automatically detected in three left (F3, C3, P3) and three right (F4, C4, P4) homologous electrodes in linked-mastoid-referenced EEG-signals filtered between 0.5 and 4 Hz (Riedner *et al.*, 2007). A peak-to-peak amplitude threshold corresponding to 75 μV was applied. Of note, this threshold was selected as it is a criterion adopted in the clinical practice for the definition of slow waves; however, similar results were also obtained using a 40 μV negative amplitude threshold. The density of slow waves in each epoch and channel was computed as described above. Finally, the average absolute inter-hemispheric difference in slow wave density was computed across pairs of homologous electrodes.

### Probabilistic origin and recruitment

Next, we evaluated whether slow waves originate with a different incidence across the two hemispheres. Thus, individual slow waves were classified as having a left-hemisphere (right-hemisphere) origin if > 75% of the origin-channels (delay = 0 ms) were located in the left (right) hemisphere. An origin asymmetry index was computed as the difference in the density (waves/min) of slow waves originating in the left vs. the right hemisphere. In addition, we defined the slow wave “*probabilistic origin*” as the probability for each channel to represent the origin of a slow wave, computed with respect to all the detected slow waves. Similarly, the “*probabilistic recruitment*” was defined as the probability for each electrode to be part of the propagation path of a slow wave.

### Statistical Analyses

For each parameter of interest, the 5 CP and the 3 NP (Ancona dataset) were compared with the control group of healthy adult individuals (HS; Lausanne dataset). Specifically, for each patient, the relative z-score and corresponding p-value were computed with respect to the distribution represented by the healthy control group. A Bonferroni correction was applied to account for multiple comparisons across tested subjects and related hypotheses. Effects were regarded as significant only when a corrected *P* < 0.05 was observed in each of the 5 CP and in none of the 3 NP (Supplementary Table S3 summarizes group- and subject-level statistics for each performed comparison). Analyses were repeated after regression-based adjustment of values to account for inter-subjects age differences. For analyses performed in individual groups (HS, CP) against the null-hypothesis of no inter-hemispheric asymmetry, a bootstrapping procedure (1,000 iterations) was applied to compute confidence intervals (bCI, α = 0.05).

### Data availability

Relevant data that support the findings of this study are available from the corresponding authors upon motivated request.

## Results

### Sleep structure

Table 1 displays the sleep macro-structure in each patient and in the healthy control group. Examined NREM epochs in CP were characterized by an increase in SWA (0.5-4Hz) and a decrease in high-frequency activity (18-35Hz) relative to wake epochs (Supplementary Figure S3), thus confirming the reliability of performed sleep scoring.

### Slow wave characteristics

Slow wave density (CP = 15.8 ± 4.2 waves/min, range 9.8-17.5; HS = 18.7 ± 4.4 waves/min, range 9.8-25.5), amplitude (CP = 62.3 ± 21.5 μV, range 50.3-102.8; HS = 50.3 ± 15.6 μV, range 32.1-97.9), slope (CP = 1.6 ± 0.5 μV/ms, range 1.4-2.5; HS = 1.1 ± 0.3 μV/ms, range 0.8-1.8) and propagation speed (CP = 2.0 ± 0.3 m/s, range 1.5-2.5; HS = 2.3 ± 0.3 m/s, range 1.8-2.9) did not differ between CP and healthy controls as a group (Figure 1). Specifically, we found no significant differences (*P*_*cor.*_ < 0.05) in slow wave density (all *P*_*unc.*_ > 0.04, |z| < 2.0491), while significant effects were observed only in CP03 for amplitude (*P*_*unc.*_ = 0.008, |z| = 3.3688), in CP03 (*P*_*unc.*_ < 0.001, |z| = 5.1634) and CP04 (*P*_*unc.*_ < 0.001, |z| = 4.1840) for slope and in CP05 (*P*_*unc.*_ = 0.002, |z| = 3.1509) for speed. Results did not change after controlling for between-subjects age differences, except for speed, in which no significant differences were found in any patients with respect to HS (all *P*_*unc.*_ > 0.01, |z| < 2.5532).

**Figure 1.**
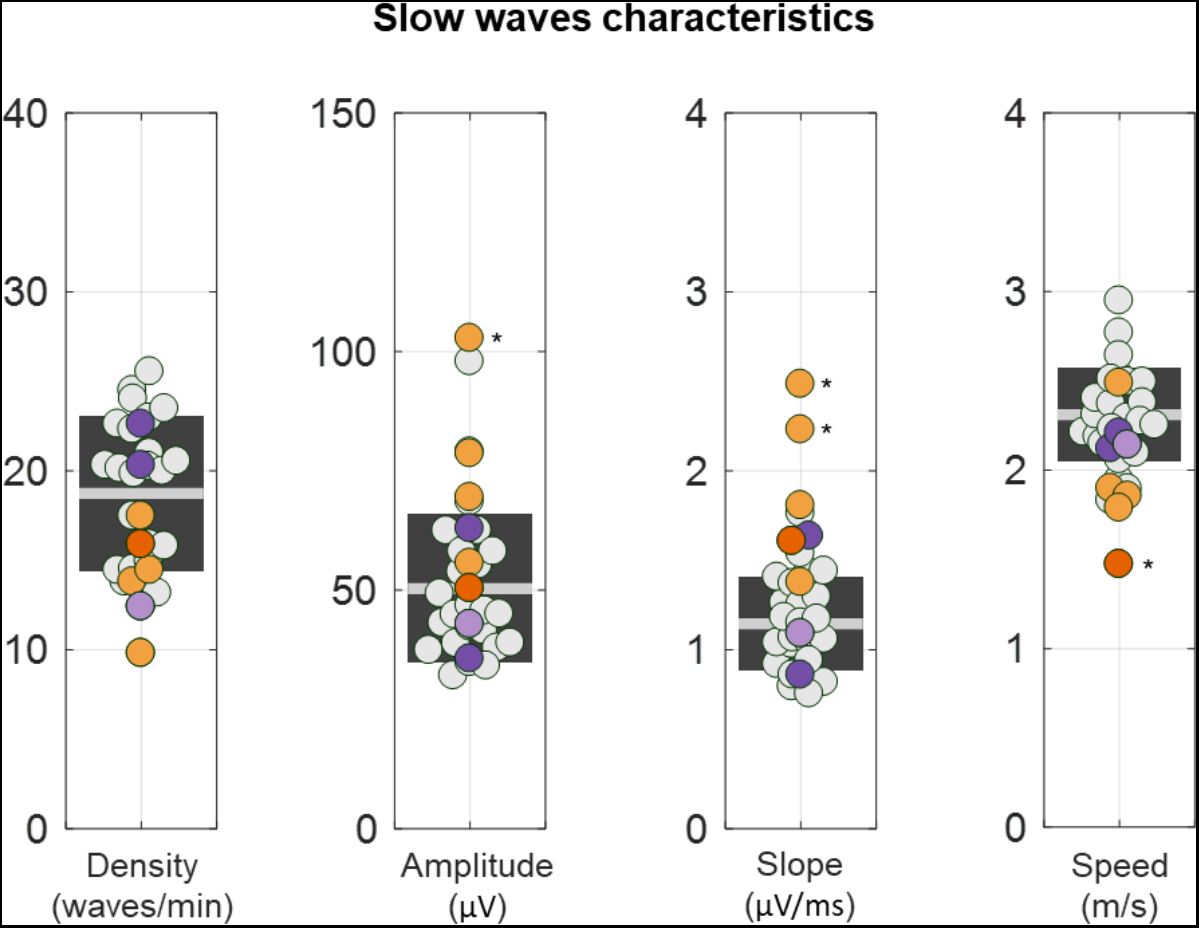
Properties of NREM slow waves. For each subject we computed slow wave density (number of waves per minute), negative amplitude (μV), descending slope (μV/ms), and propagation speed (m/s). CP are represented with orange dots (CP05 = dark orange), NP with purple dots (NP03 = light purple), and HS with light-gray dots. The light-gray horizontal line represents the mean for the HS, while the dark-gray box represents 1 SD around the mean. Values observed in the 8 patients (CP and NP) were compared with the 24 HS. * P_cor._ < 0.05.

### Slow wave involvement

Visual inspection of EEG-traces suggested that most sleep slow waves of CP may present an asymmetric scalp distribution (Figure 2).

**Figure 2.**
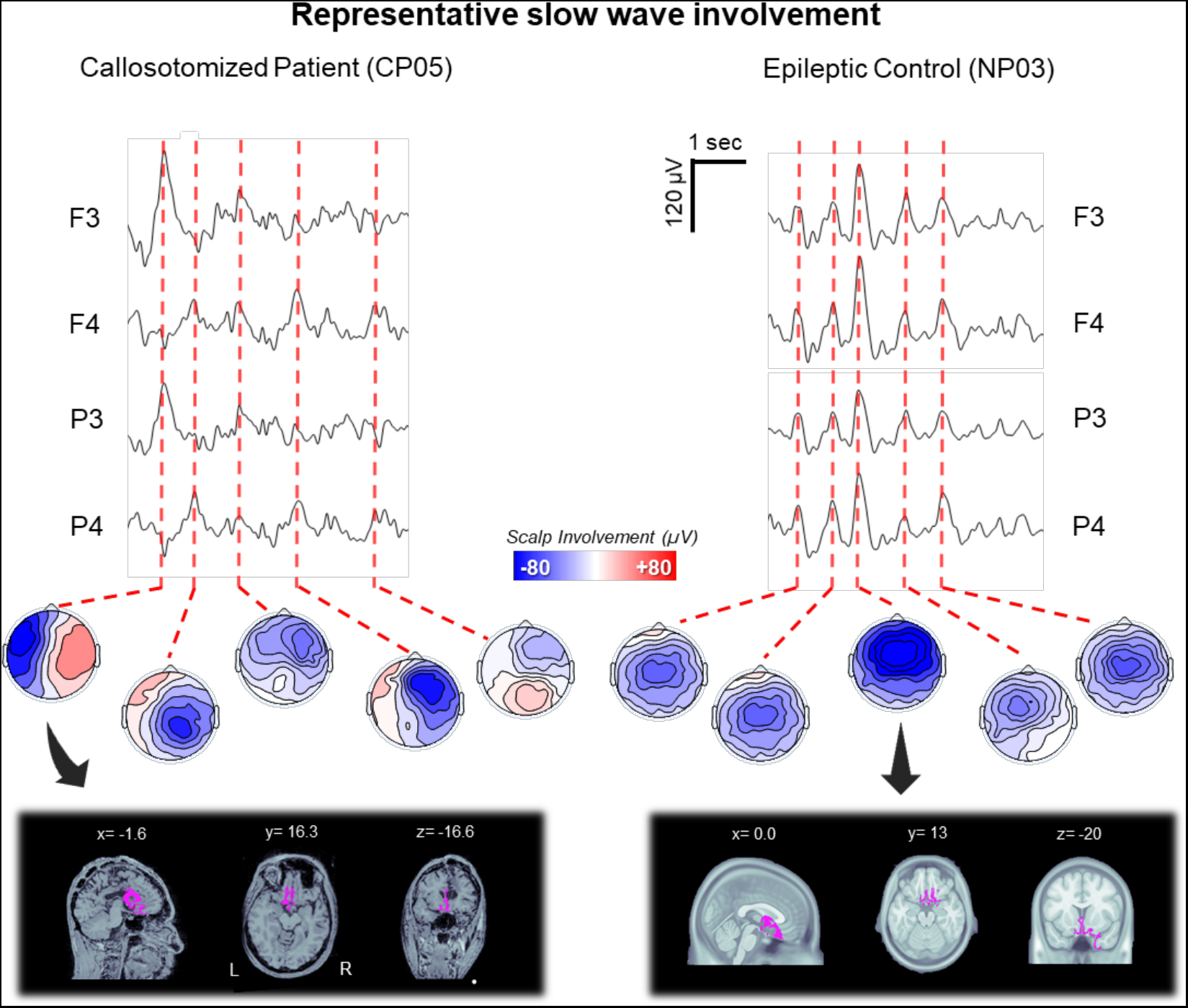
Representative slow wave involvement patterns in a callosotomized (CP05) and a non-callosotomized (NP03) epileptic patient. The top panel shows representative NREM-sleep EEG-traces (5-s) for two left (F3, P3) and two right (F4, P4) channels and the relative scalp involvement associated with exemplar slow waves. The bottom panel shows the source-reconstructed signal distributions for two representative slow waves. The source modeling has been performed using BrainStorm. A symmetric Boundary Element Method (BEM) was used to define the forward model, while the inverse matrix was computed using the standardized low-resolution brain electromagnetic tomography (sLORETA) constraint. The cortical maps are thresholded at 80% of the maximum signal amplitude.

This observation was quantitatively confirmed through a PCA-based comparison of slow wave involvement across groups (Figure 3). In fact, in HS the 95% of the variance related to scalp slow wave involvement was explained by three PCs, with maxima located in the centro-frontal area (73.1 ± 7.0%, range 57.4-85.3, of the total variance explained by the first three components), anterior or posterior areas (19.7 ± 5.7%, range 9.4-34.6) and left or right hemispheres (7.2 ± 3.1%, range 2.6-15.4), respectively. Similar values were obtained in the NP-group, with percentages corresponding to 73.9 ± 7.1% (range 67.9-81.8), 14.9 ± 3.3% (range 11.2-17.2), 11.2 ± 4.4% (range 7.0-15.8), respectively. On the other hand, in the CP-group we observed a significant increase in the variance explained by the third (left/right) component (39.0 ± 9.5%, range 29.8-53.6; *P*_*cor.*_ < 0.05, |z| > 7.3503; Bonferroni correction based on the number of tested subjects and PCs), at the expenses of the other two symmetrical components (43.7 ± 14.1%, range 26.8-61.7, for the centro-frontal component and 17.2 ± 9.6%, range 8.5-32.0, for the anterior/posterior component). In particular, the variance explained by the first component was significantly decreased in four (CP01, CP03, CP04, CP05) out of five CP.

**Figure 3.**
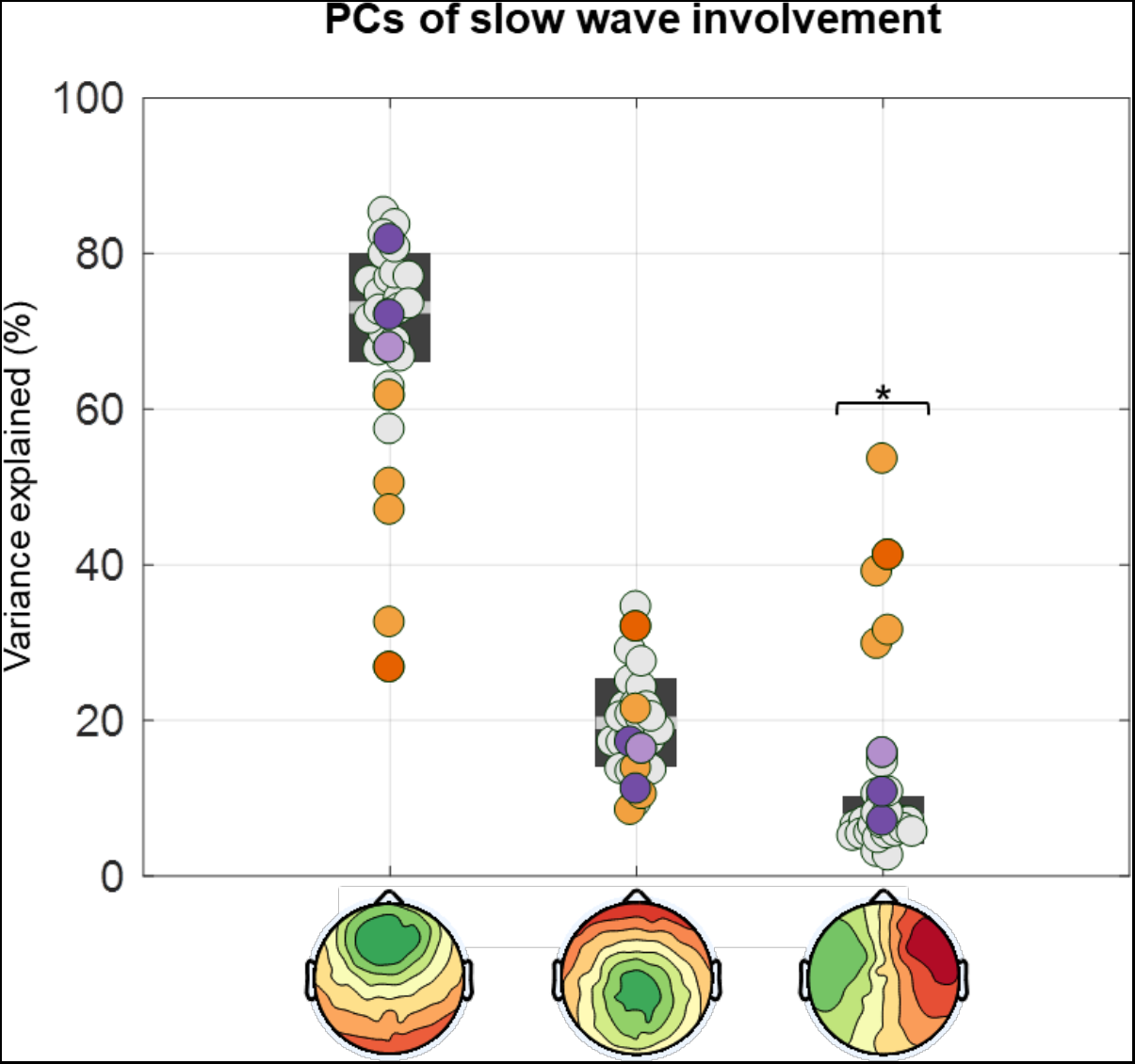
PCA-based analysis of slow wave involvement. The involvement distribution (mean EEG-signal calculated across all electrodes in a 40-ms window centered on the wave peak; μV) of all slow waves was entered in a principal component analysis (PCA). The plot shows the variance explained by each of the three PCs in all subjects. CP are represented with orange dots (CP05 = dark orange), NP with purple dots (NP03 = light purple), and HS with light-gray dots. * P_cor._ < 0.05.

### Cross-hemispheric propagation of slow waves

Next, we investigated whether alterations in the scalp distribution of slow waves in callosotomized patients could be explained by a lack of cross-hemispheric propagation of individual slow waves (Figure 4).

**Figure 4.**
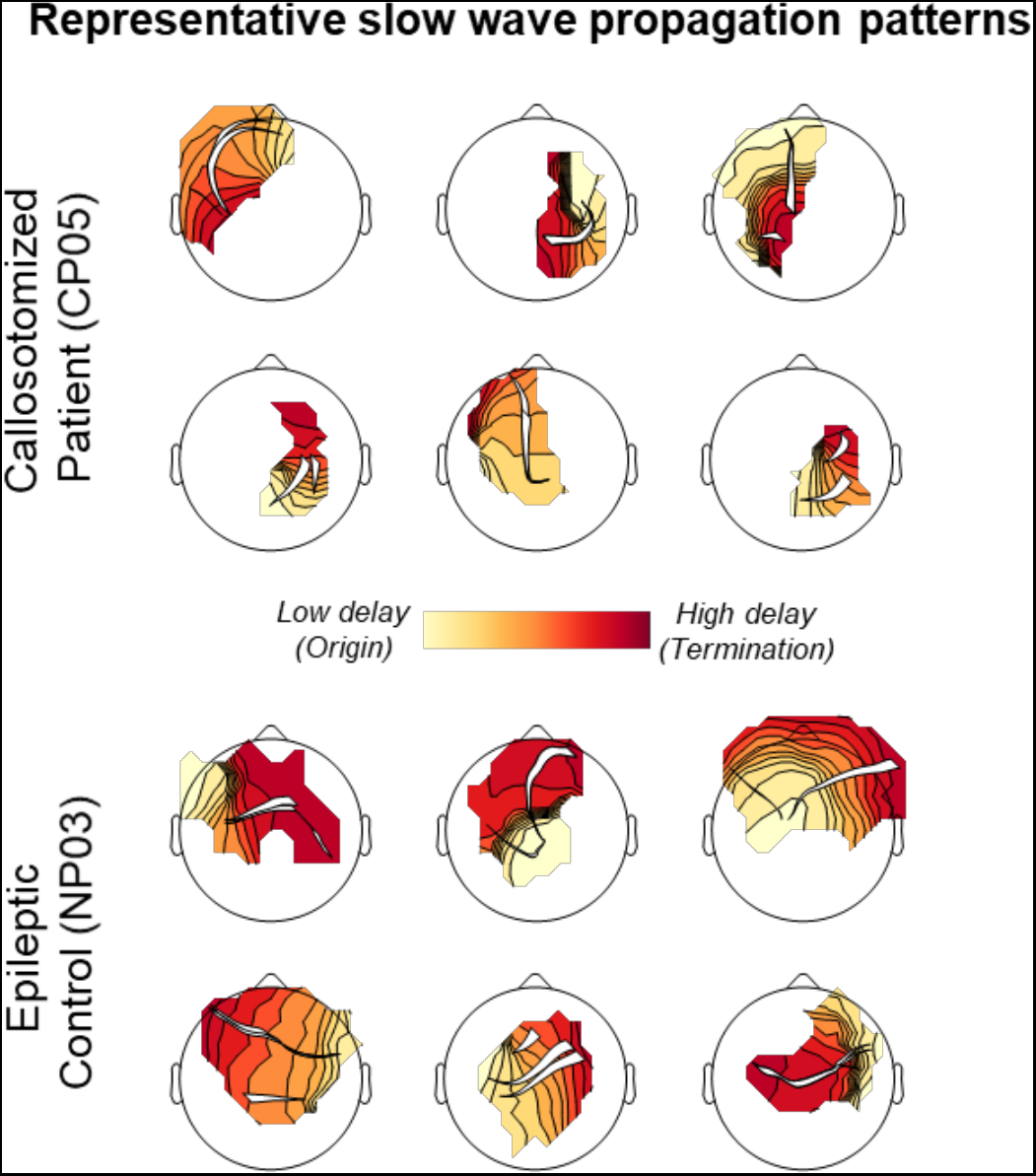
Representative slow wave propagation patterns. Traveling delay-maps and relative propagation streamlines of six representative slow waves of CP05 and NP03. In CP slow waves tended to remain confined to the origin hemisphere, while cross-hemispheric propagation was common in NP and HS.

The percentage of slow waves presenting a cross-hemispheric propagation was significantly reduced in CP (37.0 ± 8.6%, range 22.4-43.6) relative to HS (63.2 ± 3.5%, range 54.6-69.0; *P*_*cor.*_ < 0.05, |z| > 5.5345; Figure 5A). Consistent, slow waves in CP showed a stronger lateralization in terms of number of channels recruited along the propagation pattern in each of the two hemispheres (CP = 21.8 ± 4.0%, range 19.7-24.8; HS = 36.8 ± 2.4%, range 33.4-43.3; *P*_*cor.*_ < 0.05, |z| > 4.9546; Figure 5B). Of note, such lateralization appeared to similarly affect all the slow waves regardless of their amplitude (*P*_*cor.*_ < 0.05, Bonferroni correction based on the number of tested subjects and amplitude percentile classes; Figure 5C). All results remained significant after controlling for between-subjects age differences.

**Figure 5.**
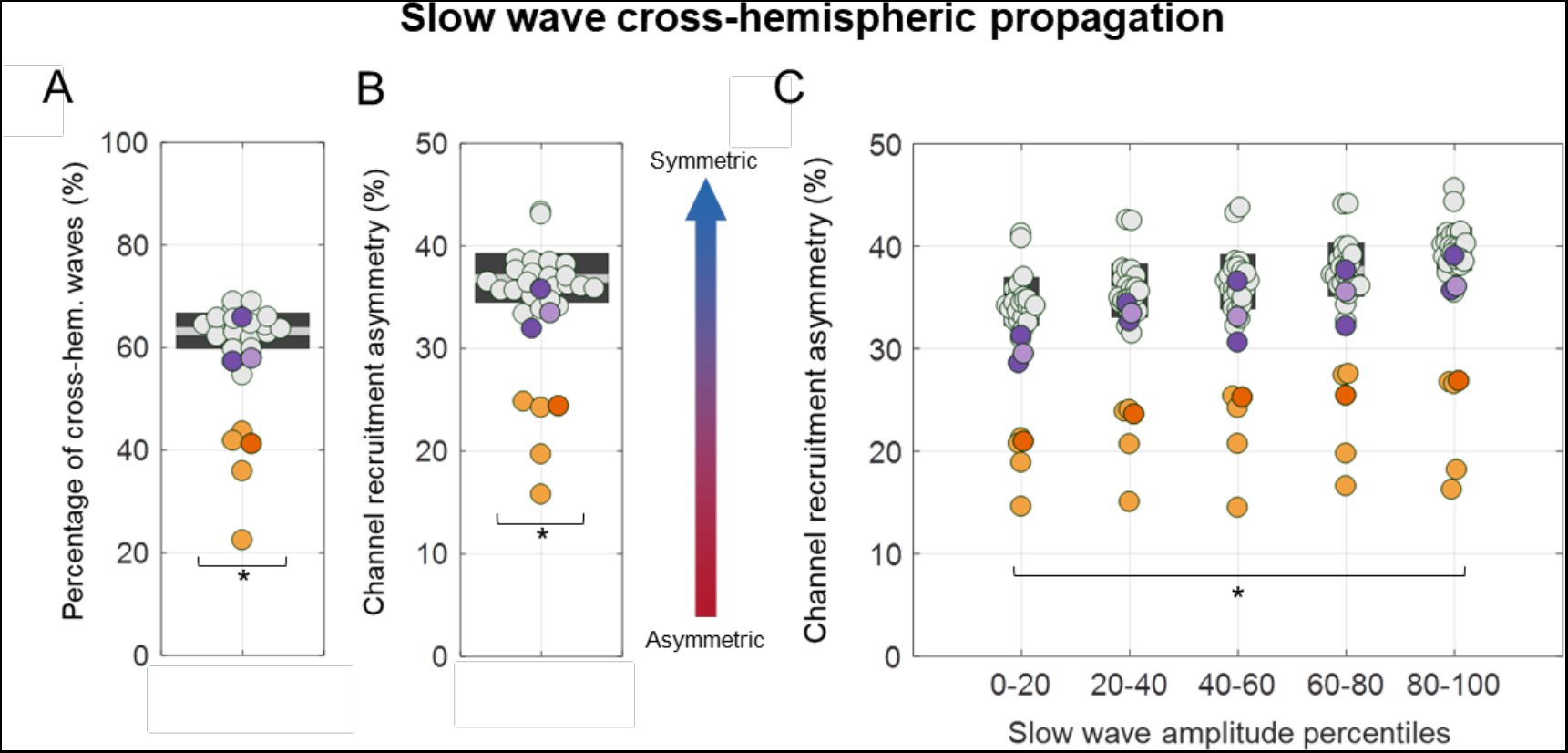
Quantitative analysis of slow wave cross-hemispheric propagation. **A)** The percentage of cross-hemispheric slow waves was computed as the number of slow waves for which at least one of the propagation streamlines passed the nasion-inion, midline axis relative to the total number of detected slow waves. **B)** The recruitment asymmetry was determined by computing the number of channels in the hemisphere with less recruited electrodes divided by the total number of recruited channels across the two hemispheres. Values close to 50% indicate a symmetric distribution, while values close to 0% indicate a unilateral wave. **C)** This second parameter was also computed for slow waves grouped into five amplitude percentile classes (0-20, 20-40, 40-60, 60-80, 80-100). CP present a significantly reduced percentage of cross-hemispheric slow waves and an increased channel recruitment asymmetry (uni-hemispheric distribution). CP are represented with orange dots (CP05 = dark orange), NP with purple dots (NP03 = light purple), and HS with light-gray dots. * P_cor._ < 0.05.

Supplementary Figure S4 shows the probabilistic channel recruitment of slow waves originating in the left or right hemisphere in each of the CP individuals and in the HS group. This qualitative representation further shows that slow waves tended to remain confined to the origin hemisphere in callosotomized but not in control subjects.

### Inter-hemispheric asymmetry in sleep depth

Last, we investigated whether the lack of strong inter-hemispheric connections could be responsible for an unbalanced sleep depth - as reflected by the generation and synchronization of sleep slow waves - across the two hemispheres. To this aim, we first tested whether CP and control HS presented an asymmetric incidence of large amplitude slow waves, characterized by peak-to-peak (negative-to-positive) amplitude greater than 75 μV. Specifically, we computed for each sleep epoch the relative difference in slow wave incidence across homologous electrodes of the two hemispheres. We found that, in both groups, many of the NREM-sleep epochs were characterized by an inter-hemispheric difference in slow wave density (Supplementary Figure S5), with a relative hemispheric dominance that varied from epoch to epoch. Overall, however, slow wave density tended to be higher in the right, relative to the left, hemisphere in both CP (−3.38 ± 0.69 waves/min, difference left-right) and HS (−0.26 ± 0.15 waves/min), although the effect reached statistical significance only in the first group (one sample t-tests against the null hypothesis of no asymmetry; HS, *P* = 0. 099, |t_23_| = 1.7197, bCIs = [−0.55, 0.01]; CP, *P* = 0.012, |t_4_| = 4.3676, bCIs = [−4.84, −2.08]). Similar results were obtained using a slow wave amplitude threshold corresponding to a negative amplitude of 40μV (HS, *P* < 0.074, |t_23_| = 1.8735, bCIs = [−0.65, −0.01]; CP, *P* = 0.017, |t_4_| = 3.9427, bCIs = [−5.39, −2.04]). Importantly, the relative inter-hemispheric asymmetry was not systematically different across CP and HS (Supplementary Table S3). However, when the absolute (|left – right|), rather than the relative (left – right) inter-hemispheric difference in slow wave incidence was considered, this parameter was significantly greater in CP (5.6 ± 3.8 waves/min) relative to HS (1.7 ± 0.5 waves/min; *P*_*cor.*_ < 0.05, |z| > 4.6552; Figure 6). Similar results were obtained for the 40μV amplitude threshold (HS = 1.9 ± 0.5 waves/min; CP = 6.2 ± 3.9 waves/min; *P*_*cor.*_ < 0.05, |z| > 5.8069). All results remained significant after controlling for between-subjects age differences.

**Figure 6.**
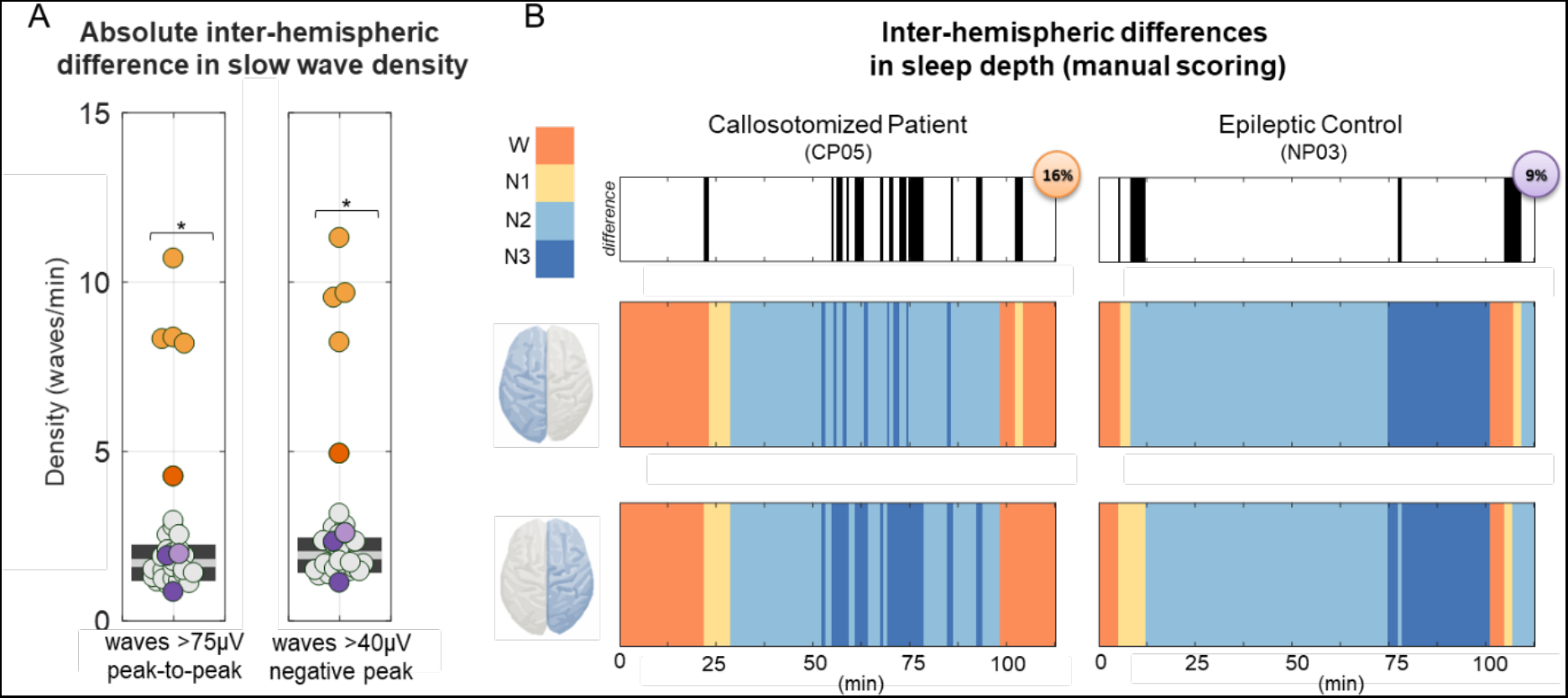
Inter-hemispheric difference in the incidence of large slow waves. **A)** Absolute inter-hemispheric difference in slow wave density. The plot on the left shows the absolute inter-hemispheric (|left-right|) difference in slow wave density. CP are represented with orange dots (CP05 = dark orange), NP with purple dots (NP03 = light purple), and HS with light-gray dots. * P_cor._ < 0.05. **B)** Sleep scoring (120 minutes; 0 = time of lights off) in a callosotomized patient (CP05) and in the non-callosotomized epileptic patient (NP03). Sleep scoring was performed separately for each hemisphere (i.e., using only electrodes of the left or right side) by an operator blind to both the identity of the subjects and the evaluated brain hemisphere. In the top panel, black sections indicate epochs for which different stages were scored across the two hemispheres. Bottom panels represent the sleep scoring for the first sleep cycle in the two patients and for each hemisphere (left top, right bottom).

In light of the above observations, we then asked whether the greater inter-hemispheric differences in slow wave incidence found in CP could be better explained by a more disproportionate slow wave generation across brain hemispheres, or simply by the lack of cross-hemispheric propagation. To this aim, we first determined the overall proportion of slow waves with a clear origin in the left or in the right hemisphere with respect to the total number of detected slow waves (Figure 7). Consistent with the above-reported results, we found that a greater percentage of waves originated in the right hemisphere in both HS (paired t-test, *P* = 0.009, |t_23_| = 2.8471, bCIs = [−4.11, −0.85]; left = 39.95 ± 2.99%, right = 42.45 ± 2.47%) and CP (paired t-test, *P* = 0.03, |t_4_| = 3.4595, bCIs = [−17.64, −5.58]; left = 37.33 ± 4.04%, right = 49.03 ± 3.59%). However, the relative and the absolute inter-hemispheric asymmetry in origin density were not significantly different across groups based on the predefined statistical criteria (Supplementary Table S3).

**Figure 7.**
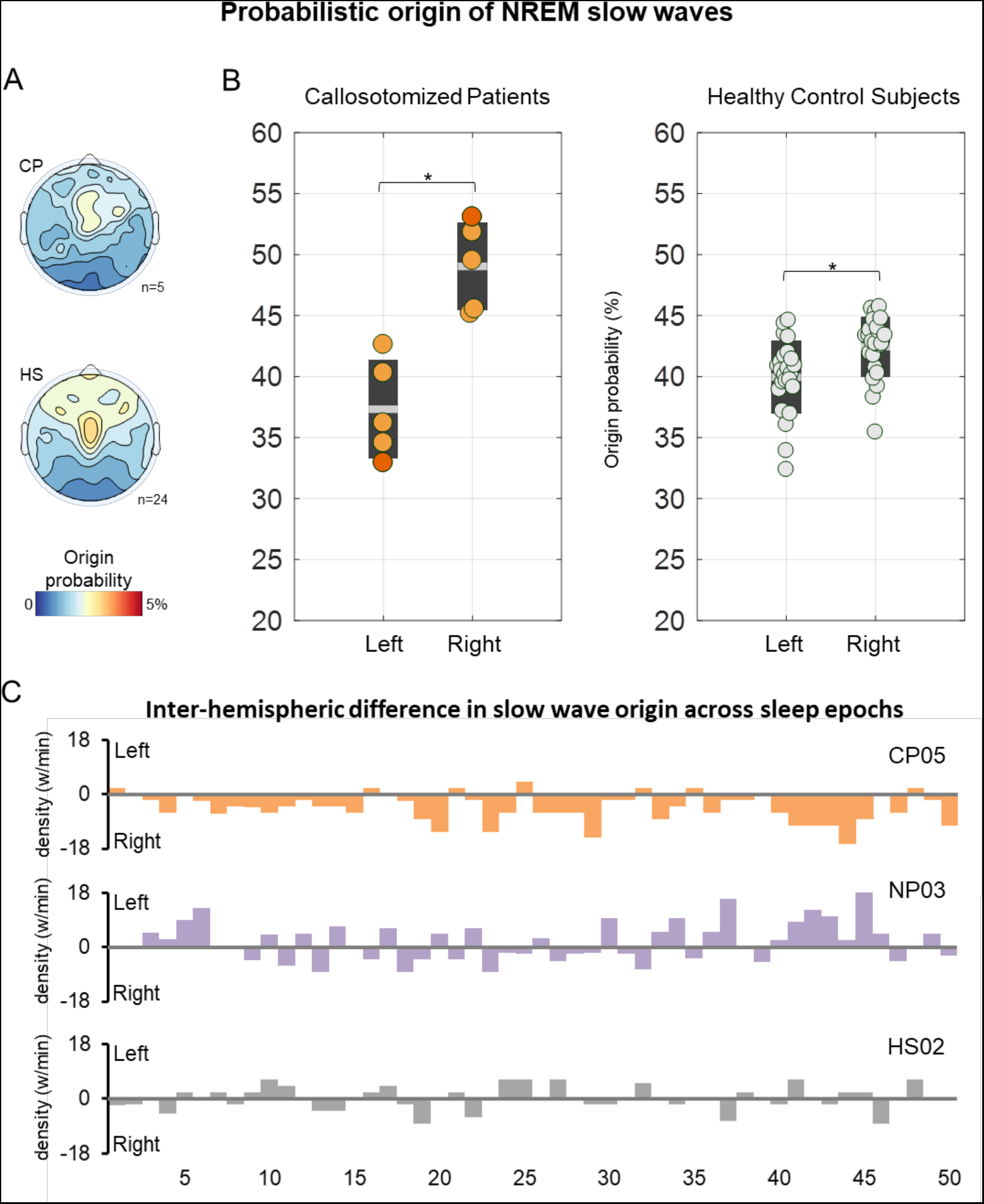
Differences in probabilistic origin across the left and right hemisphere. **A)** Topographic map of probabilistic slow wave origin in CP (top) and HS (bottom). Similar distributions, with maxima in central-lateral and anterior areas were found in both CS and HS. **B)** A higher proportion of slow waves originated in the right vs. left hemisphere in both CP and HS. CP are represented with orange dots (CP05 = dark orange), NP with purple dots (NP03 = light purple) and HS with light-gray dots. * P_cor._ < 0.05. **C)** Inter-hemispheric difference in slow wave origin density across the first 50 epochs of NREM-sleep in three representative subjects (CP05, NP03, HS02). For each epoch, the bar represents the relative difference in origin density between the left and the right hemisphere (left – right). Thus, a positive value indicates higher density in the left hemisphere, and vice-versa.

## Discussion

The slow waves of NREM-sleep have been shown to spread across brain areas in scalp hd-EEG recordings of healthy human individuals (Massimini *et al.*, 2004; Murphy *et al.*, 2009). While this macro-scale cortical traveling has been thought to be mediated by cortico-cortical white matter connections, to date only indirect, correlational evidence has supported this assumption (Buchmann *et al.*, 2011; Piantoni *et al.*, 2013). Furthermore, findings reported in the literature are contradictory, likely because of methodological discrepancies and limitations. In the present study we show that a complete resection of the corpus callosum (CC), which contains the main bundle of inter-hemispheric white matter fibers, is associated with an increased incidence of uni-hemispheric slow waves, reflecting a decrease in cross-hemispheric propagation. Interestingly, our results also demonstrate that slow waves originate more often in the right relative to the left hemisphere and that this asymmetry is not significantly affected by the resection of the CC.

### The corpus callosum is responsible for the cross-hemispheric propagation of sleep slow waves

Here we demonstrate that the cross-hemispheric propagation of NREM slow waves largely depends on the integrity of callosal white matter tracts. Indeed, while in healthy adult subjects more than the 60% of all slow waves showed a clear cross-hemispheric propagation, in callosotomized patients more than the 60% of them remained confined within the cerebral hemisphere in which they originated. These results are in line with previous correlational evidence indicating a direct relationship between parameters reflecting slow wave synchronization and the (micro)structure of the anterior CC (Buchmann *et al.*, 2011; Piantoni *et al.*, 2013; but see also Sanchez *et al.*, 2019). More in general, they provide support to the hypothesized relationship between patterns of slow wave propagation and structural cortico-cortical connectivity (Murphy *et al.*, 2009; Kurth *et al.*, 2017; Schoch *et al.*, 2018). In this respect, our findings are somewhat in contrast with recent work showing a positive correlation between indices reflecting white matter damage and slow wave synchronization efficiency in TBI patients (Sanchez *et al.*, 2019). These discrepancies may be explained by the different methodological approach, as the indices investigated in the previous work (slow wave amplitude/slope) only indirectly reflect slow wave cortical synchronization/propagation across brain areas. Another possibility is that a damage-related “disconnection” in TBI patients may enhance the cortical propensity to locally generate and synchronize slow-wave-like events, as shown in animal models of cortical deafferentation (Timofeev *et al.*, 2000; Topolnik *et al.*, 2003).

The slow-wave cortical traveling has been suggested to have a direct role in organizing information processing and plasticity in cortical networks through the local modulation of spindles and high-frequency activity (Cox *et al.*, 2014). Thus, the alteration of cross-hemispheric propagation may lead to an alteration of plasticity-related processes requiring the interaction and coordination of activity across the two brain hemispheres. It should be noted, however, that the resection of the CC did not completely eliminate cross-hemispheric slow wave propagation. This may imply the existence of additional propagation pathways, potentially including cortico-subcortico-cortical loops (e.g., Timofeev and Steriade, 1996), or the involvement of direct subcortico-cortical synchronization mechanisms (Siclari *et al.*, 2014; Bernardi *et al.*, 2018). Of note, a visual inspection of the structural MRI-scans also revealed that the anterior and posterior commissures were relatively spared in all the CP individuals. However, given their small size, these structures alone are unlikely to support the observed residual cross-hemispheric propagation (Mancuso *et al.*, 2019). On the other hand, the relatively low spatial resolution of the scalp EEG and the effects of volume conduction may be expected to determine an apparent cross-hemispheric propagation, especially for slow waves originating and/or traveling spatially close to the brain midline. Future studies exploiting approaches with higher spatial resolution, such as intracranial EEG, will be necessary to clarify this issue.

### The resection of the corpus callosum is not sufficient for the manifestation of uni-hemispheric sleep

Our study showed that, during NREM-sleep, large slow waves are often asymmetrically distributed across the two hemispheres in healthy adult individuals. Interestingly, the absolute degree of inter-hemispheric asymmetry is significantly increased in callosotomized patients. Based on the standard definition of sleep stages, this particular condition could lead to apparent differences in sleep depth across the two brain hemispheres. Such an asymmetry could be explained either by a change in the number of slow waves originated in the two hemispheres, or simply by the loss of cross-hemispheric propagation after callosotomy. However, since the cortical distribution of slow wave origins was not systematically and significantly affected by the resection of the CC, the observed asymmetry could be entirely explained by the reduced cross-hemispheric slow wave propagation. This observation is in line with previous evidence suggesting that the lack or resection of inter-hemispheric connections is not sufficient for the manifestation of uni-hemispheric sleep (Berlucchi, 1966; Montplaisir *et al.*, 1990; Nielsen *et al.*, 1993), as naturally seen in some animal species, such as birds and cetaceans (Rattenborg *et al.*, 2000; Mascetti, 2016). On the other hand, the absence (as in birds) or small size (as in cetaceans) (Tarpley and Ridgway, 1994) of the CC may prevent the cross-hemispheric spreading of sleep slow waves, and thus represent one fundamental prerequisite for uni-hemispheric sleep.

### Slow waves originate more often in the right than in the left hemisphere

Present results revealed that during NREM-sleep, slow waves tend to originate more often in the right than in the left hemisphere in healthy adult subjects as well as in callosotomized patients, although the relative hemispheric predominance also varies from epoch to epoch. A similar inter-hemispheric difference in SWA during NREM-sleep has been reported in some previous investigations (e.g., Goldstein *et al.*, 1972; Sekimoto *et al.*, 2000, 2007). Interestingly, our observation of a similar lateralization in patients who underwent callosal resection implies that such slow wave lateralization does not depend on competitive regulatory mechanisms acting across the two hemispheres. Why then does the right hemisphere generate more slow waves than the left one during NREM-sleep? In light of the homeostatic mechanisms that regulate SWA (Borbely and Achermann, 1999) and of the known differences in hemispheric functional specialization (Karolis *et al.*, 2019), the right hemisphere may develop a stronger function- and use-dependent “sleep need” during wakefulness that translates into higher slow wave activity during subsequent sleep. However, this possibility is at odds with previous findings indicating a stronger rebound in SWA within the left hemisphere following extended wakefulness, relative to baseline sleep conditions (Achermann *et al.*, 2001; Ferrara *et al.*, 2002; Vyazovskiy *et al.*, 2002). Of note, a recent work showed that the first night of sleep in a new environment may be associated with an increased sleep-depth asymmetry, with the left hemisphere operating as a “night watch” (Tamaki *et al.*, 2016). This observation raises the interesting possibility of a constitutional difference in the arousal-related, bottom-up control of sleep in the two hemispheres. One could speculate that a “deeper sleep” of the right hemisphere, highly involved in attentional control, may enable a relative disengagement from environmental stimuli (Bareham *et al.*, 2014), while a “more awake” left hemisphere could facilitate the recognition of potentially relevant communicative stimuli that are especially important in social animals (Legendre *et al.*, 2019). More specific studies will be required to directly put these hypotheses into test.

### Limitations

The main limitation of this study is the relatively small sample size. However, it should be emphasized that patients who underwent complete callosotomy represent an exceptionally rare population (Fabri *et al.*, 2017). Furthermore, to overcome potential limitations related to the sample size, we performed evaluations at single subject level and applied strict criteria for the definition of “significant” group differences. Another potential limitation is that all the epileptic patients (CP01-CP05 and NP03) presented alterations in the background EEG activity caused by the underlying pathological condition. All the signals have been carefully inspected to discard segments containing non-physiological activity. Though, it is still possible for some slow-wave-like epileptic events to have been included in our analyses. In addition, the use of medications, including anti-epileptic and hypnotic drugs (Supplementary Table S2), may also have affected recorded EEG-signals. Although we cannot completely exclude the influence of these factors on our analyses, all results were consistent across a heterogeneous sample of callosotomized patients with distinct underlying conditions, comorbidities and pharmacological therapies. Moreover, the non-callosotomized patients, including an epileptic subject, who were studied under similar conditions did not show the same pattern of slow wave differences observed in callosotomized patients as compared to the healthy adult control group.

## Conclusions

This study systematically investigated the origin, distribution and traveling of sleep slow waves in complete split-brain patients. To the best of our knowledge, our results are the first demonstration that the resection of inter-hemispheric connections significantly limits the cross-hemispheric propagation of sleep slow waves without affecting the relative distribution of slow wave origins across the two hemispheres. These findings also provide further support to previous assumptions regarding the dependence of slow waves on cortico-cortical connections for their macro-scale spreading. In light of previous evidence indicating that slow waves may modulate, throughout their propagation, spindle and high-frequency activity potentially related to plastic processes, our results indicate that callosotomy may significantly affect these sleep-dependent mechanisms. On a different perspective, our findings also demonstrate that the loss of inter-hemispheric connections in adult life is not sufficient, *per-se*, to allow the appearance of uni-hemispheric sleep in humans, thus implying that in animals showing this particular behavioral state additional functional and/or anatomical mechanisms may play a pivotal role.

## Abbreviations

CC: Corpus callosum
CP: Callosotomized patients
Hd-EEG: High-density Electroencephalography
HS: Healthy subjects
NP: Non-callosotomized patients
SWA: Slow wave activity
TBI: Traumatic brain injury

## Acknowledgements

The authors wish to thank all the patients, their families and the volunteers who took part in the study and the personnel of the Neurological Unit of the Ancona Regional Hospital for their help and assistance with data acquisition. The authors wish to thank also Gabriella Venanzi for the precious support she offered to the patients and their families.

## Author Contributions

### Conceptualization

G. Bernardi, M. Bellesi, M. Silvestrini, G. Tononi, P. Pietrini, F. Siclari, E. Ricciardi, M. Fabri, G. Avvenuti;

### Investigation

G. Avvenuti, G. Bernardi, M. Bellesi, J. Cataldi, S. Lattanzi, G. Polonara;

### Data Curation

G. Bernardi, G. Avvenuti;

### Formal Analysis

G. Avvenuti, G. Bernardi, G. Handjaras, M. Betta, L.S. Imperatori;

### Methodology

B.A. Riedner, G. Bernardi;

### Visualization

G. Avvenuti, G. Bernardi;

### Resources

P. Pietrini, E. Ricciardi, G. Tononi, B.A. Riedner, F. Siclari, M. Silvestrini, M. Fabri, G. Polonara;

### Supervision

G. Bernardi, M. Bellesi;

### Writing – original draft

G. Bernardi, G. Avvenuti, M. Bellesi;

### Writing – review & editing

All authors.

