## Supplementary Material for "Integrity of corpus callosum is essential for the cross-hemispheric propagation of sleep slow waves: a high-density EEG study in split-brain patients"

* *Equal contribution*

#### Table S1

| STUDY | MODEL | SUBJECTS | METRIC | STATISTICS | RESULTS |
| --- | --- | --- | --- | --- | --- |
| Kuks et al., 1987 | AgCC | 3 infants (age ≤ 5 months) | Coherence | Comparison with normal infants | Reduced inter-hemispheric EEG-coherence in AgCC |
| Montplaisir et al., 1990 | ReCC | 1 epileptic adult with anterior ReCC; 1 epileptic adult with posterior ReCC | Coherence | Comparison before-after surgery | Reduced inter-hemispheric EEG-coherence after ReCC |
| Nielsen et al., 1993 | AgCC | 4 young adult subjects | Coherence | Comparison with a group of healthy adult subjects | Reduced inter-hemispheric EEG-coherence in AgCC |
| Corsi-Cabrera et al., 2006 | ReCC | 1 epileptic adult with complete ReCC; 1 epileptic adult with anterior ReCC | Correlation Spectra | Comparison before-after surgery and against a group of control healthy subjects | Attenuation of inter-hemispheric correlation after ReCC but preserved correlation increase during sleep relative to wakefulness |

*Table S1. Previous human sleep studies in subjects with agenesis of the corpus callosum and in epileptic patients who underwent partial or complete resection of the corpus callosum. AgCC = Agenesis of the corpus callosum; ReCC = Resection of the corpus callosum.*

**Table S2**

| | DIAGNOSED PATHOLOGY | CURRENT MEDICATIONS |
|---|---|---|
| **CP01** | Lennox-Gastaut Syndrome | Carbamazepine, Phenytoin sodium, Phenobarbital |
| **CP02** | Drug-resistant epilepsy | Carbamazepine, Levetiracetam, Sodium valproate |
| **CP03** | Early Infantile Epileptic Encephalopathy | Levosulpiride, Oxcarbazepine, Phenobarbital, Risperidone |
| **CP04** | Lennox-Gastaut Syndrome | Clobazam, Lacosamide, Phenobarbital, Rosuvastatin, Vigabatrin |
| **CP05** | Drug-resistant epilepsy | Carbamazepine, Clonazepam, Diazepam, Omeprazole, Phenobarbital |
| **NP01** | Generalized Anxiety Disorder | None |
| **NP02** | Lumbar spinal stenosis | Cholecalciferol, Esomeprazole, Lisinopril, Mometasone |
| **NP03** | Epilepsy (viral meningoencephalitis in infancy) | Enalapril+Lercanidipine, Lacosamide, Oxcarbazepine, Sodium valproate, Ursodeoxycholic acid |

*Table S2. Clinical diagnosis and medications of studied patients. CP = callosotomized patients; NP = non-callosotomized patients.*

### Table S3

| | | HEALTHY SUBJECTS (N=24) | | | | | NON-CALL. PATIENTS | | | CALLOSOTOMIZED PATIENTS | | | | |
|---|---|---|---|---|---|---|---|---|---|---|---|---|---|---|
| ANALYSIS | PARAMETER | KS TEST | MEAN | SD | PRC. 2.5 | PRC. 97.5 | NP01 | NP02 | NP03 | CP01 | CP02 | CP03 | CP04 | CP05 |
| Slow wave properties | Density | 0.350 | 18.7 | 4.4 | 10.1 | 25.4 | 20.3 | 22.6 | 12.4 | 13.8 | 14.4 | 17.5 | 9.8 | 15.9 |
| | Amplitude | 0.410 | 50.3 | 15.6 | 32.3 | 96.0 | 62.9 | 35.6 | 42.8 | 55.6 | 78.6 | **102.8** | 69.4 | 50.3 |
| | Slope | 0.350 | 1.1 | 0.3 | 0.8 | 1.7 | 1.6 | 0.9 | 1.1 | 1.4 | **1.8** | **2.5** | **2.2** | 1.6 |
| | Speed | 0.991 | 2.3 | 0.3 | 1.8 | 2.9 | 2.1 | 2.2 | 2.1 | 1.9 | 2.5 | 1.9 | **1.8** | **1.5** |
| * Involv. PCA | FrontoCentral | 0.993 | 73.1 | 7.0 | 57.8 | 85.1 | 81.8 | 72.1 | 67.9 | **32.6** | 61.7 | **50.4** | **47.0** | **26.8** |
| | Anter./Poster. | 0.744 | 19.7 | 5.7 | 9.7 | 34.0 | 11.2 | 17.2 | 16.3 | 13.9 | **8.5** | 10.5 | 21.4 | 32.0 |
| | Left/Right | 0.119 | 7.2 | 3.1 | 2.7 | 15.3 | 7.0 | 10.8 | **15.8** | **53.6** | **29.8** | **39.1** | **31.5** | **41.2** |
| Tra.v. Asy. | Crosshem. Pr. | 0.830 | 63.2 | 3.5 | 54.9 | 69.0 | 65.8 | 57.3 | 57.8 | **35.9** | **43.6** | **22.4** | **41.8** | **41.2** |
| | Asymmetry | 0.652 | 36.8 | 2.4 | 43.3 | 33.4 | 35.8 | **31.9** | 33.4 | **19.7** | **24.8** | **15.8** | **24.2** | **24.4** |
| * Asymmetry amplitude percentiles | Prc. 0-20 | 0.414 | 34.6 | 2.4 | 31.1 | 41.2 | 31.3 | **28.6** | **29.5** | **21.2** | **20.7** | **14.6** | **18.9** | **20.9** |
| | Prc, 20-40 | 0.560 | 35.7 | 2.7 | 31.5 | 42.5 | 34.4 | 32.6 | 33.4 | **20.7** | **23.8** | **15.0** | **24.0** | **23.6** |
| | Prc, 40-60 | 0.622 | 36.5 | 2.7 | 32.3 | 43.7 | 36.5 | **30.6** | 33.1 | **20.7** | **25.3** | **14.5** | **24.2** | **25.2** |
| | Prc, 60-80 | 0.865 | 37.7 | 2.6 | 32.8 | 44.1 | 37.6 | **32.2** | 35.5 | **19.7** | **27.4** | **16.6** | **27.5** | **25.4** |
| | Prc, 80-100 | 0.696 | 39.8 | 2.1 | 35.6 | 45.5 | 39.0 | 35.7 | 36.0 | **16.2** | **26.7** | **18.2** | **26.5** | **26.8** |
| Relat. Hem. Diff. | Density 75μV | 0.869 | -0.3 | 0.7 | -2.5 | 1.0 | 0.2 | 0.2 | -0.5 | **-5.4** | **-3.4** | -1.8 | **-4.6** | -1.5 |
| | Density 40μV | 0.839 | -0.3 | 0.8 | -2.6 | 1.1 | 0.1 | 0.2 | -1.0 | **-5.7** | **-4.1** | -1.5 | **-5.7** | -1.6 |
| | Origins | 0.908 | -0.5 | 0.7 | -2.1 | 0.9 | **1.0** | -0.2 | **2.5** | **-3.4** | **-3.1** | -1.4 | -0.3 | **-3.8** |
| Absol. Hem. Diff. | Density 75μV | 0.382 | 1.7 | 0.5 | 1.1 | 2.9 | 1.9 | **0.8** | 2.0 | **8.3** | **8.4** | **10.7** | **8.2** | **4.3** |
| | Density 40μV | 0.192 | 1.9 | 0.5 | 1.3 | 3.1 | 2.3 | **1.1** | 2.6 | **8.2** | **9.5** | **11.3** | **9.7** | **4.9** |
| | Origins | 0.967 | 4.4 | 0.6 | 3.0 | 5.5 | 4.1 | 5.0 | **6.3** | **7.4** | **6.8** | **7.1** | 3.7 | 5.5 |

*Table S3. Summary of statistics related to comparisons between patients and healthy subjects. The first two columns indicate the analyses of interest. Columns three to seven include descriptive statistics for the healthy subjects (HS) group: p-value of the Kolmogorov–Smirnov test for data normality (KS Test), group level mean (Mean), standard deviation of the mean (SD), 2.5 (Prc. 2.5) and 97.5 (Prc. 97.5) percentiles of the distribution. Columns from eight to ten show the values of the parameter of interest observed in each of the three non-callosotomized patients (NP01-NP03). Columns from ten to thirteen show the values of the parameter of interest observed in each of the five callosotomized patients (CP01-CP03). Bold text indicates values that fall off the 2.5-97.5 percentiles range (α < 0.05). Shaded green cells mark values that are significantly different from those of the HS group after Bonferroni correction. The correction was applied based on the number of tested subjects (N=8). Are exceptions the analyses marked with a *, for which the correction also took into account the number of related parameters that were tested.*

**Figure S1**

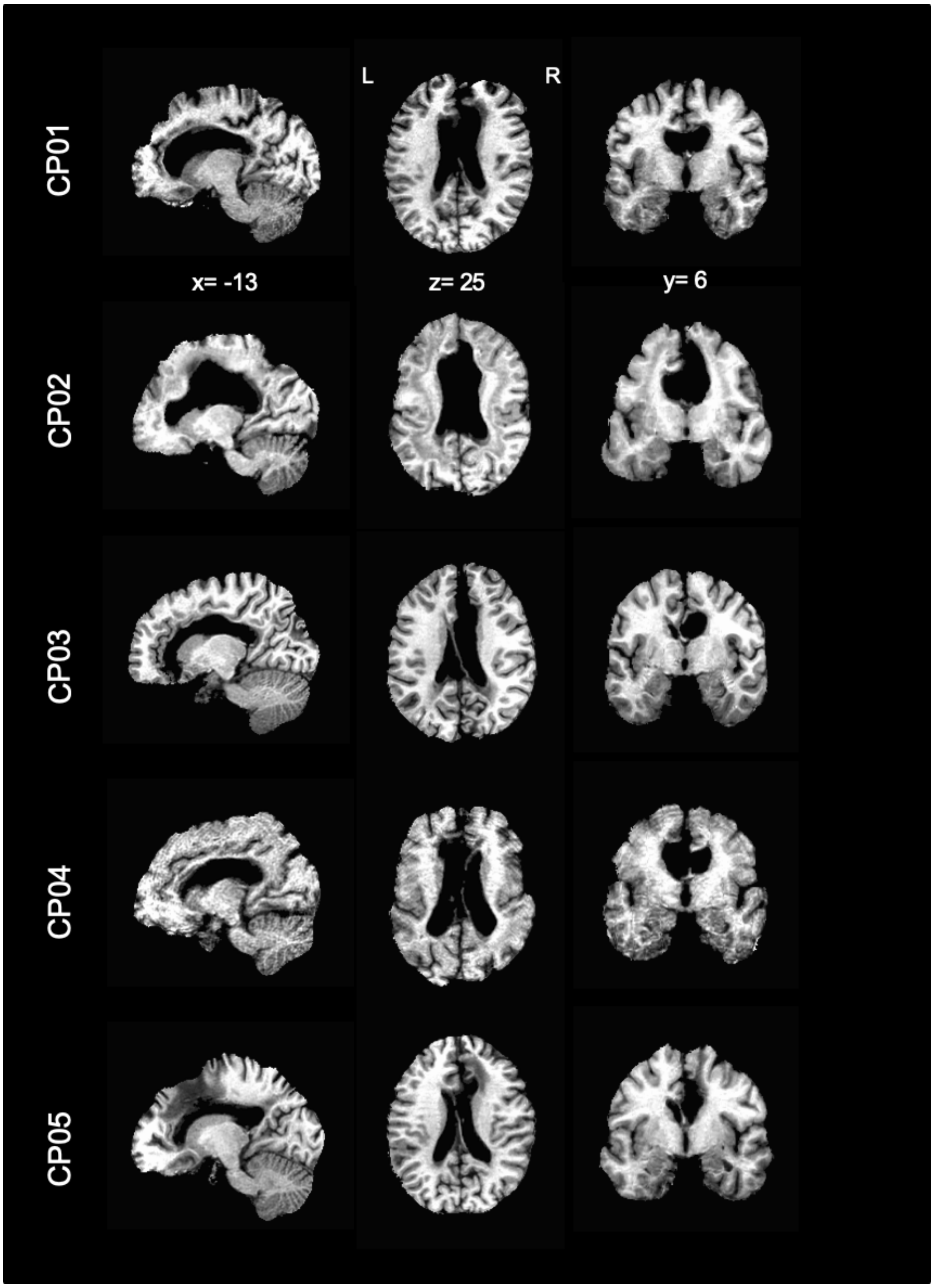


***Figure S1. Anatomical MRI images of callosotomized patients.*** *For each patient (CP01-CP05) sagittal, axial and coronal MRI images are shown in MNI space. It is possible to appreciate the complete absence of the corpus callosum in all cases.*

#### Figure S2

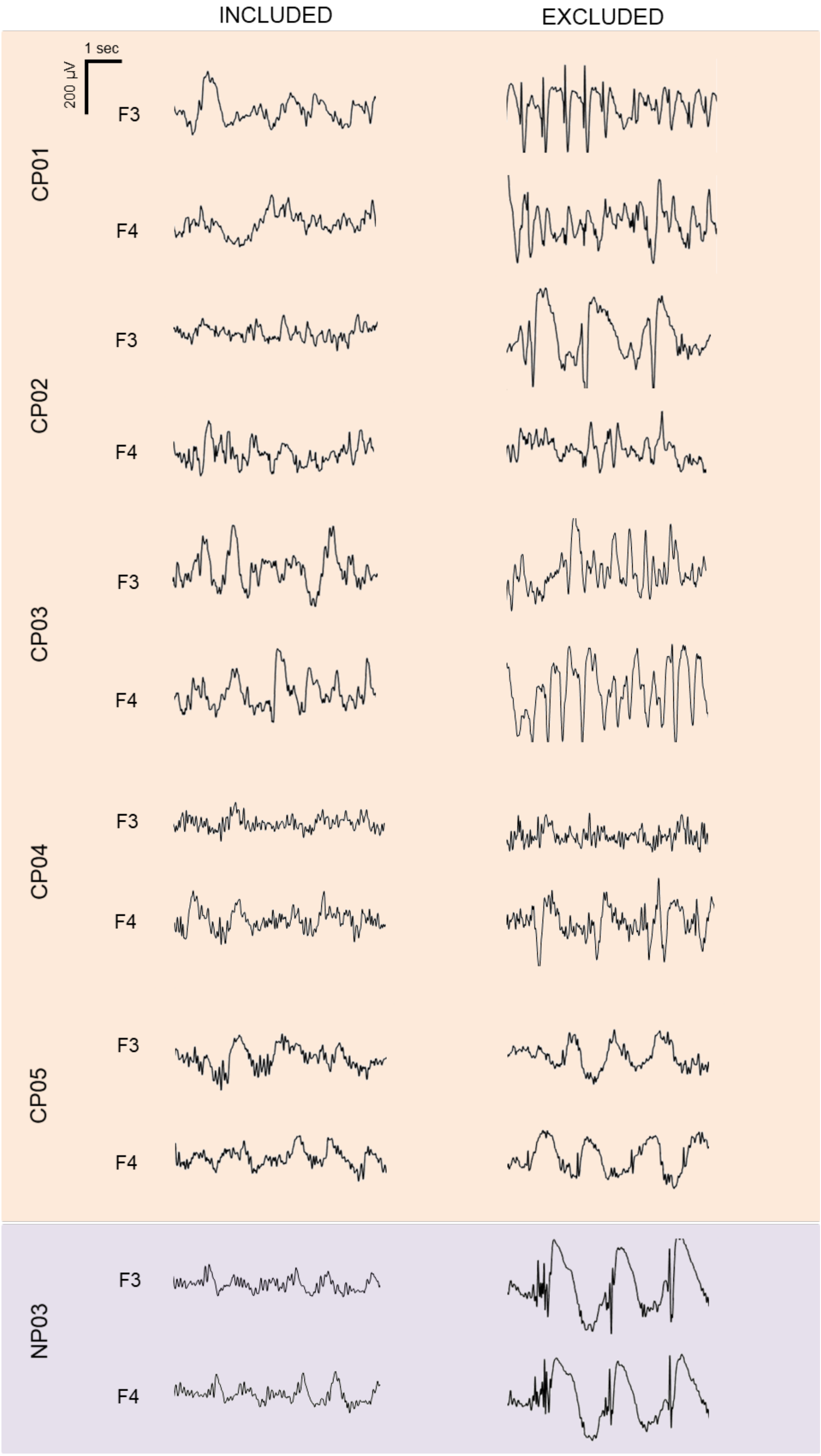


**Figure S2.** ***Examples of included and excluded data segments in epileptic patients.*** *For each subject representative EEG-traces (0.5-25 Hz) corresponding to included (left) and excluded (right) data segments are shown for one frontal left (F3) and one frontal right (F4) electrode.*

**Figure S3**

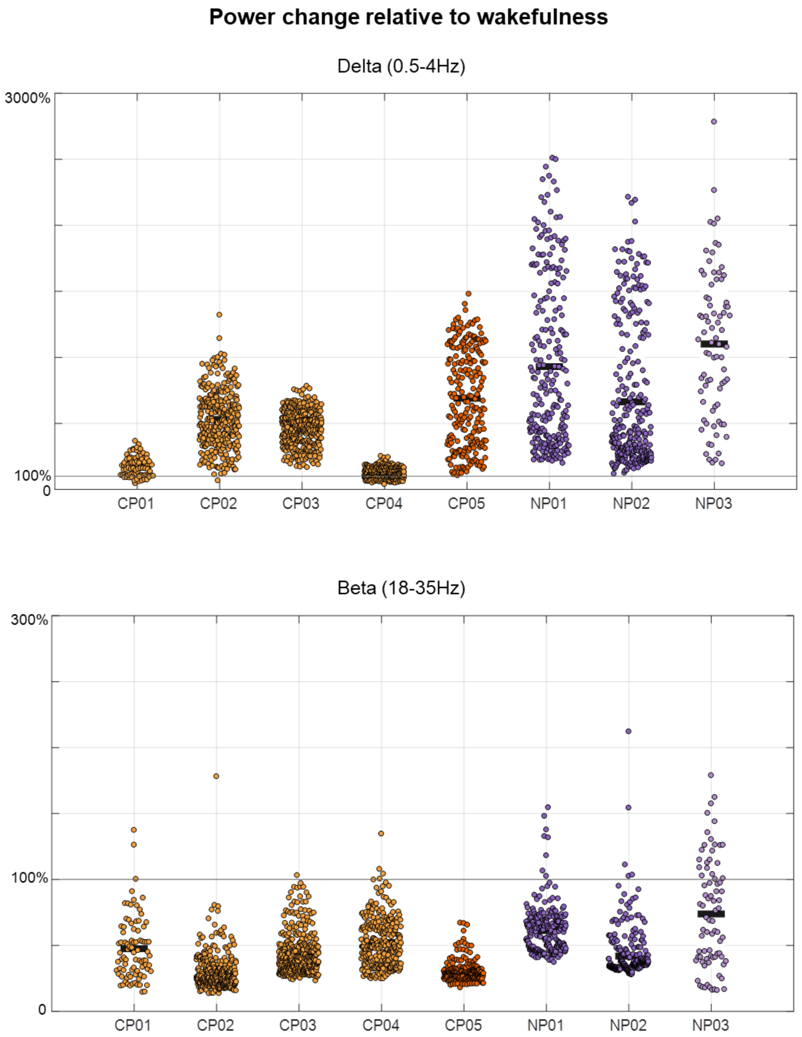


***Figure S3. EEG power changes relative to wakefulness.*** *The sleep stage classification in four of the five callosotomized patients (CP01-CP04) and of the epileptic non-callosotomized control patient (NP; NP03) was made difficult by the presence of altered patterns of brain activity. However, the reliability of the sleep scoring procedure is supported by the direct comparison of EEG power between epochs scored as NREM-sleep and eyes-closed wake recordings collected prior to sleep, which showed an increase in SWA (0.5-4 Hz) and a decrease in high-frequency activity (beta, 18-35 Hz) in all CP (CP01-CP05) and NP (NP01-NP03) subjects. Here 100% corresponds to the signal power in wakefulness. The SWA increase was overall smaller in CP relative to NP. CP = callosotomized patients; NP = non-callosotomized patients. CP are represented with orange dots (CP05 = dark orange), NP with purple dots (NP03 = light purple).*

#### Figure S4

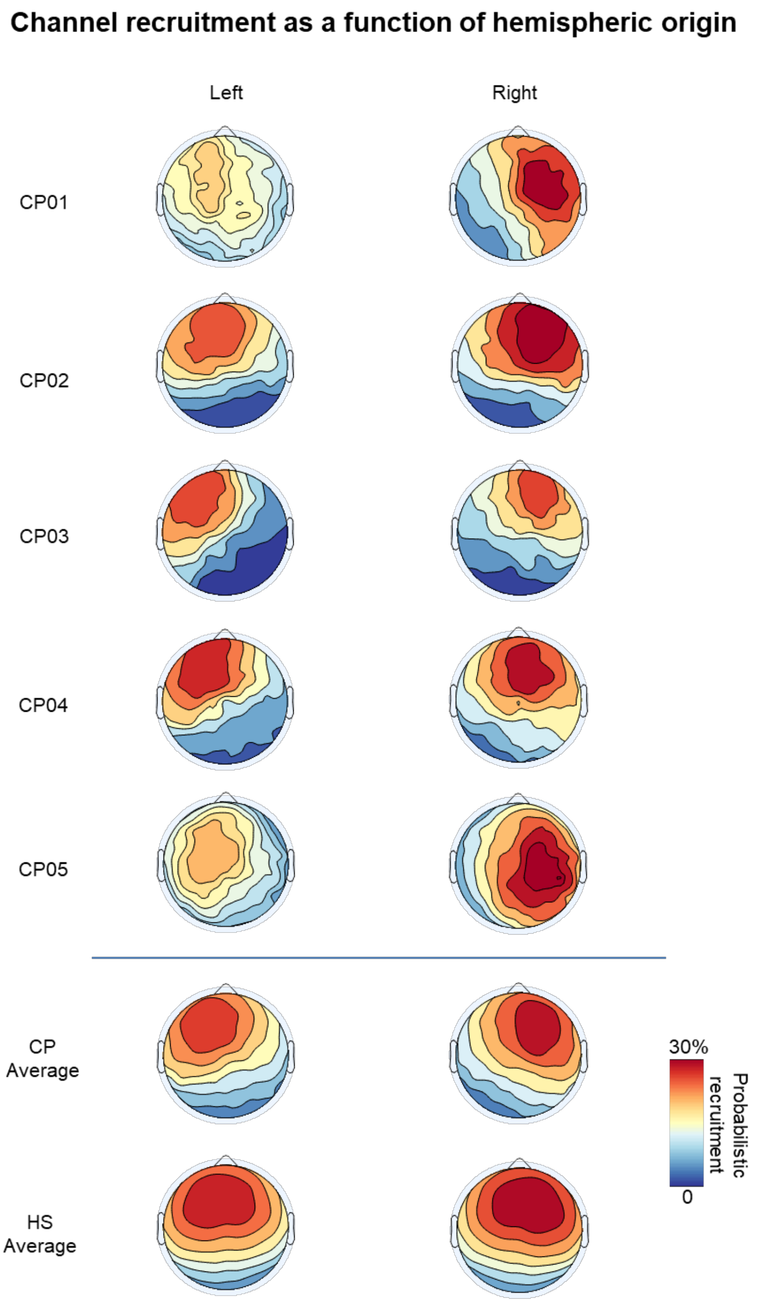


***Figure S4. Scalp probabilistic recruitment as function of hemispheric origin.*** *The top panel shows, for each of the CP, the probabilistic recruitment (probability of channel recruitment in a slow wave) for slow waves originating in the left (left-column) or right (right-column) hemisphere. The probabilistic recruitment is expressed as a percentage with respect to the total number of detected slow waves, regardless of their origin site. The bottom panel shows the average probabilistic recruitment for CP and HS. It is evident that the involvement tended to be more symmetrical in HS with respect to CP. Moreover, this analysis suggested a relatively stronger recruitment of the right (vs. left) hemisphere in both CP and HS.*

**Figure S5**

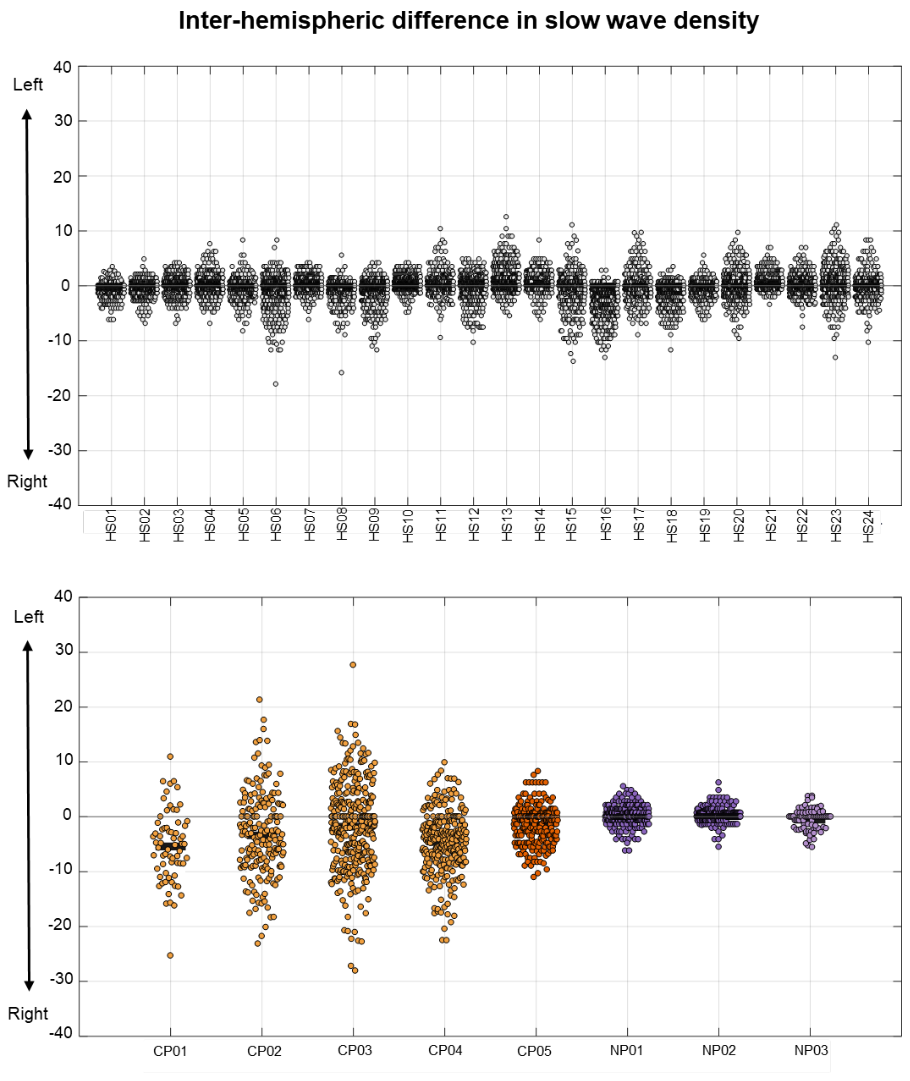


***Figure S5. Inter-hemispheric difference in slow wave density****. Difference in the mean slow wave density (waves/min) across three left (F3, C3, P3) and three right (F4, C4, P4) channels. A peak-to-peak amplitude threshold corresponding to 75 µV was applied to minimize spurious cross-hemispheric detection caused by simple volume conduction. Each dot represents a different NREM-sleep epoch. The top plot represents each of the HS (HS01-HS24), while the bottom plot shows the CP (CP01-CP05) and the NP (NP01-NP03). Lower (negative) values indicate a higher number of slow waves detected in the right hemisphere. In both CP and HS groups there was a tendency toward a higher slow wave density in the right relative to the left hemisphere.*
